# TDKC (Target Distilled K-mer Classifier): Ultrafast and Memory-Efficient Sequence Classification for Target Pathogen Diagnostics

**DOI:** 10.64898/2026.06.05.730319

**Authors:** Seungmo Lee, Vivek Agarwal, William O’Brien, Eleazar Eskin

## Abstract

Metagenomic sequencing can identify pathogens from clinical samples without prior knowledge of the causative agent. Yet, as sequencing workflows scale to process thousands of multiplexed samples simultaneously, classifying these samples against massive reference databases creates a significant computational bottleneck. Furthermore, large-scale applications such as screening public sequence repositories remain computationally challenging.

Existing metagenomic classifiers are designed for full-taxon classification, where the goal is to identify all organisms in a sample. However, many diagnostic applications focus on detecting a specific set of clinically relevant pathogens. This constraint can be exploited to significantly lower computational costs.

Here we present TDKC (**T**arget **D**istilled **K**-mer **C**lassifier), a method for targeted metagenomic classification. TDKC constructs a compact index by distilling target-specific k-mers from a full-taxon reference database. When classifying clinical samples, TDKC uses 16.9–33.6***×*** less memory and is 5.1–34.7***×*** faster than per-read full-taxon and targeted classifiers (Kraken2, Centrifuger, CLARK), while maintaining high sensitivity and low false positive rates. Against the sketch-based profiler Sylph, TDKC remains 3.8***×*** faster and uses 8.7***×*** less memory. TDKC also supports per-k-mer accession tracking across over 3 million source accessions for downstream subtype analysis, and domain-level detection of bacteria, archaea, and viruses. By reducing the index to only the pathogens of interest, TDKC makes targeted pathogen detection feasible at scale.

## Background

As the COVID-19 pandemic emphasized, accurate and timely identification of pathogens is important for both clinical care and public health management of infectious diseases. Widely used clinical viral diagnostics, such as PCR panels and amplicon-based sequencing, are highly accurate but can only detect pathogens for which assays have been specifically designed. When a novel or unexpected pathogen emerges, these assays cannot detect it until new primers or probes are developed and validated[1, 2]. Clinical metagenomics addresses this limitation by applying whole-genome or whole-transcriptome sequencing directly to clinical specimens, enabling identification of any pathogen present without prior assumptions about the causative agent. This makes metagenomics a viable “day-one” diagnostic for future pandemics[3–5].

An important step in clinical metagenomics is matching sequenced reads against a database of known pathogen genomes to identify which organisms are present in a sample. There has been substantial progress in accurate methods for this task[6], broadly referred to as metagenomic classification. This task can be divided into two categories: (1) full-taxon classification, where the goal is to identify all organisms present, and (2) targeted classification, where the goal is to detect a predefined set of organisms of interest. Most methodological progress has focused on the full-taxon setting, which typically depends on large reference databases such as RefSeq[7] and the NCBI Nucleotide Database[8]. They have been growing rapidly; for instance, the number of observed species in RefSeq has doubled approximately every three years[9]. As these databases grow, so does the computational cost of classifying sequences against them.

Growing databases is only part of the problem. Modern high-throughput sequencing platforms can generate massive volumes of data in a single run — the Illumina NextSeq 2000, for example, produces up to 540 Gb (gigabases)[10] — and workflows often process thousands of samples through multiplex library preparation. To handle this amount of data, multiple classifier instances must be run concurrently, creating a significant throughput bottleneck in computation[11]. A similar problem arises in large-scale applications such as screening public sequence repositories[12, 13] or metagenomic surveillance of wastewater[14].

To address these challenges, alignment-free classifiers have become the dominant approach for large-scale metagenomics, sacrificing a small amount of accuracy for substantial speedups over alignment-based tools like BWA[15] and Minimap2[16]. Prior work has taken three broad strategies to scale alignment-free classification. The first applies lossless compression to the full index: Centrifuge[17] and Centrifuger[18] use Burrows-Wheeler Transform (BWT)[19] and FM-index[20] schemes to compress the reference, but backward search over the BWT is inherently sequential and bottlenecks query throughput. The second uses k-mer sketching: Sylph[21] and Sourmash[22] use MinHash-based representations to improve scalability, but these approaches come at the cost of sensitivity, particularly for low-abundance or closely related organisms. The third partitions the database and streams it from disk: Kun-peng[23] and the low-memory mode of KrakenUniq[24, 25] lower peak memory at the cost of repeated I/O and index reloading.

All of these methods are designed for the same task: full-taxon classification. Clinical diagnostics, however, falls into the second category: targeted classification. Rather than characterizing the full microbial composition of a sample, the goal is to detect a defined set of clinically relevant pathogens[3]. In this context, the use of a full-taxon database is computationally inefficient, since most of the database does not contribute to the final decision. Yet standard approaches still require loading and querying the full-taxon database, introducing unnecessary overhead that scales with a rapidly growing reference database and the number of samples to be processed.

Here, we introduce TDKC (Target Distilled K-mer Classifier), a fast and memory-efficient method designed specifically for targeted metagenomic classification. TDKC distills a compact target-specific index from a full-taxon reference. The full reference is used during index construction to verify which k-mers are unique to targets, after which only the distilled k-mers are retained. To preserve taxonomic granularity, TDKC adopts the same minimizer-based logic and least common ancestor (LCA)-resolved taxonomy as Kraken2[26], an approach widely shown to provide a good balance between sensitivity and computational performance. This allows TDKC to remain capable of distinguishing closely related strains. We demonstrate that TDKC reduces memory by 16.9–33.6× and classification runtime by 5.1–34.7× relative to per-read baseline classifiers while maintaining high sensitivity and low false positive (FP) rates on contrived validation samples.

## Results

### Methods overview

Figure 1 illustrates the TDKC workflow. TDKC distills target-specific k-mers from a full-taxon database, meaning the subset of k-mers that belong to target taxa under full-taxon LCA resolution, to perform highly efficient and accurate read classification in metagenomic samples. TDKC’s fundamental goal is to run targeted classification much faster and more memory-efficiently while achieving comparable accuracy to full-taxon classification, for which we use Kraken2 as a classifier. This is not achievable using only target reference genomes, as doing so results in many false-positive classifications (Figure 2a). Instead, TDKC processes the identical set of reference sequences as the full-taxon database during index construction, utilizing both target and non-target genomes to build the distilled index. TDKC starts by finding target-relevant clades to extract target k-mers, then undergoes a two-stage process to resolve taxonomic labels and remove any ambiguous k-mers that would otherwise cause FPs. In the first stage, each target k-mer is assigned a taxID by computing the LCA. In the second stage, referred to as the “challenge phase”, we process all non-target k-mers and check for membership in the target k-mer set; any matches are ejected. This bypasses expensive LCA computation, allowing TDKC to build a target index significantly faster. Using this distilled index, users can query their metagenomic samples to identify which target pathogens are present. Furthermore, TDKC supports optional per-k-mer accession tracking (TDKC-A), mapping each k-mer to its source accessions across more than 3 million target accessions. TDKC-A also outputs a per-sample strain report that, for each source accession, counts the distinct minimizers mapped to it and the subset unique to that accession. These per-accession counts can support downstream subtype analysis to distinguish pathogen variants. TDKC also supports domain-level detection (TDKC-D) covering bacteria, archaea, and viruses for taxonomic profiling. These steps are discussed in further detail in the “Methods” section.

**Fig. 1.**
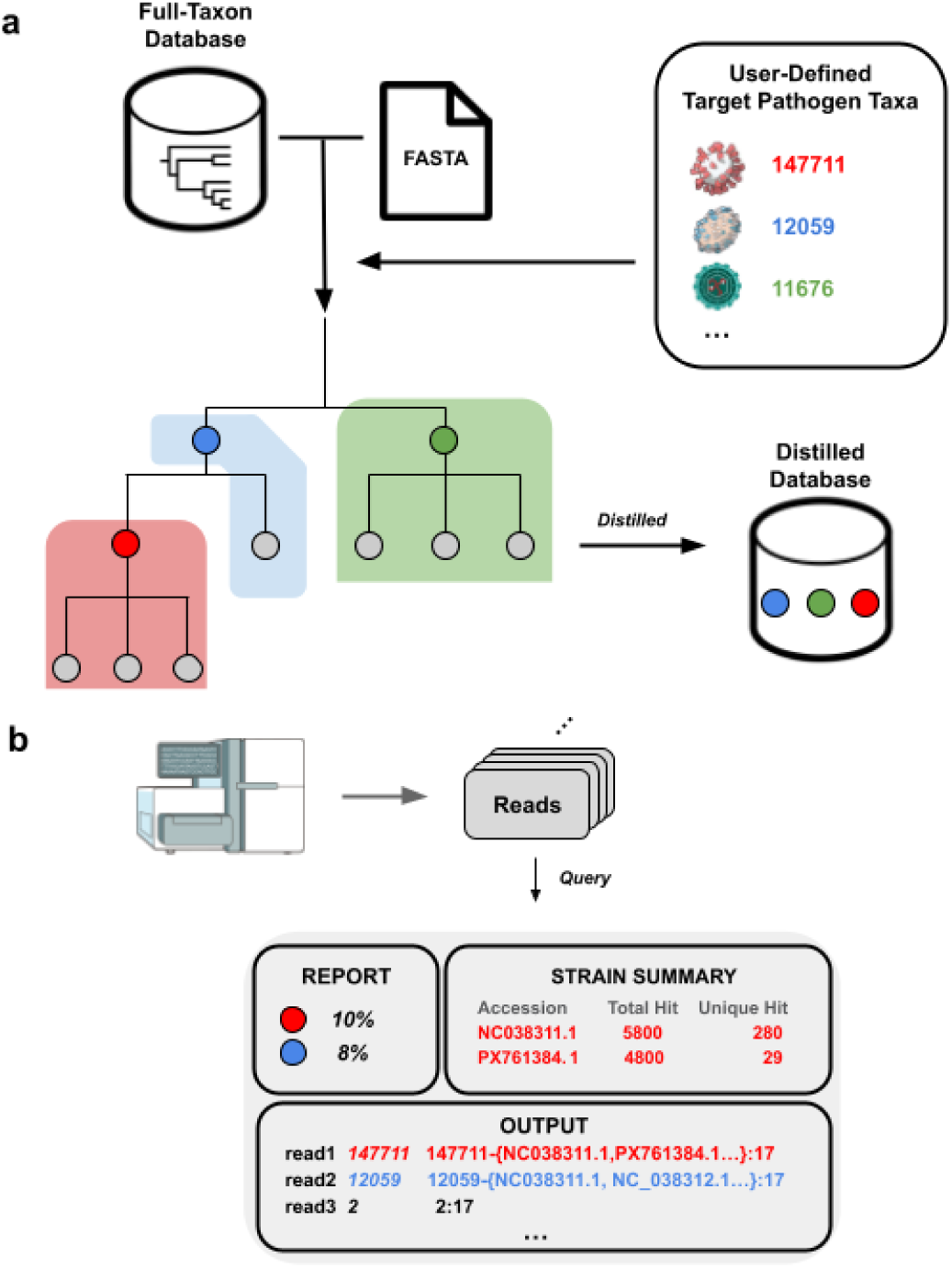
Overview of TDKC workflow. (a) Index construction. Given a full-taxon reference FASTA, the NCBI taxonomy, and a user-defined list of target pathogen taxonomy IDs (taxID), TDKC extracts target-relevant minimizers, resolves their taxonomy via LCA, and removes any minimizers shared with non-target sequences to produce a compact distilled index. (b) Query and reporting. Reads are classified using the distilled index, with optional per-k-mer accession tracking for subtype analysis and optional domain-level detection.

**Fig. 2.**
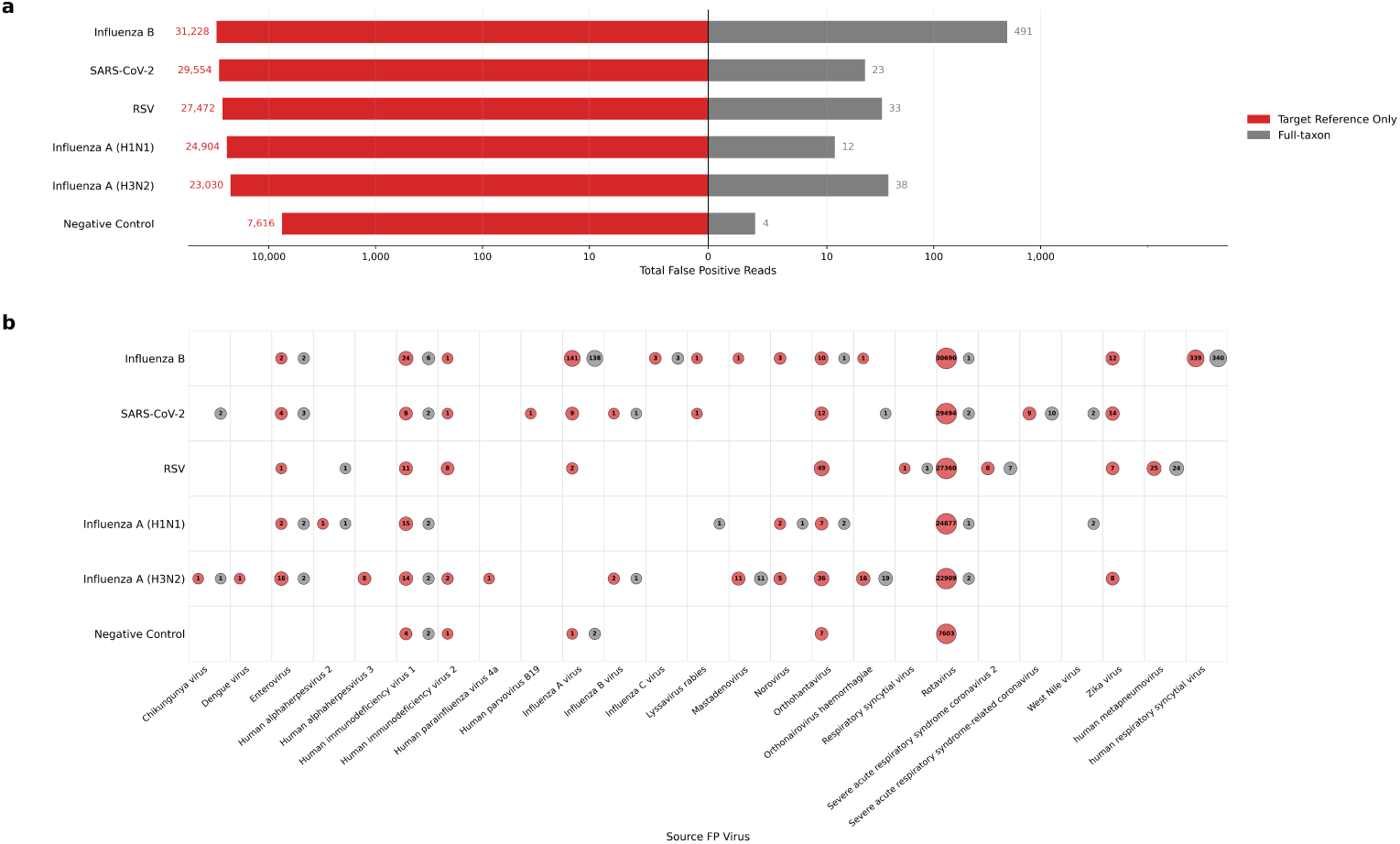
FP read counts on contrived LOD samples between databases built with target reference only and full-taxon reference sequences. Two Kraken2 databases are compared: one built from target reference sequences alone, and the other from the full-taxon reference sequences. The latter is equivalent to the full-taxon database used in the main results. The same evaluation framework as the main results is used. (a) Total FP reads per spike-in virus, aggregated across all samples, with target reference only on the left (red) and full-taxon on the right (gray). The x-axis is symlog-scaled. (b) FP reads broken down by virus they were classified as. Rows are spike-in, and columns are source FP viruses. Circle areas are proportional to the log FP read count. Empty cells indicate no FP reads under either database.

### TDKC outperforms existing classifiers in memory and speed

We benchmarked TDKC against four baseline classifiers: Kraken2, Centrifuger, and Sylph for full-taxon and CLARK[27] for targeted, using 96 multiplexed clinical nasopharyngeal swab samples (see Methods for details). All indexes were loaded once per job and samples were processed sequentially using 32 threads. All FASTQ files were gzipped.

TDKC was 5.1× faster and used 33.6× less memory than full-taxon Kraken2 (179s vs. 911s; 2.8 GB vs. 94.2 GB) (Figure 3). Against CLARK, which indexes the same target accessions as TDKC, TDKC was 34.7× faster (vs. 6,218s) and used 16.9× less memory (vs. 47.2 GB). Against Centrifuger, TDKC was 10.5× faster (vs. 1,881s) and used 30.0× less memory (vs. 83.9 GB). Against Sylph, which performs species-level abundance profiling through k-mer sketching rather than per-read classification, TDKC was 3.8× faster (vs. 680s) and used 8.7× less memory (vs. 24.4 GB), though this comparison spans different computational tasks.

**Fig. 3.**
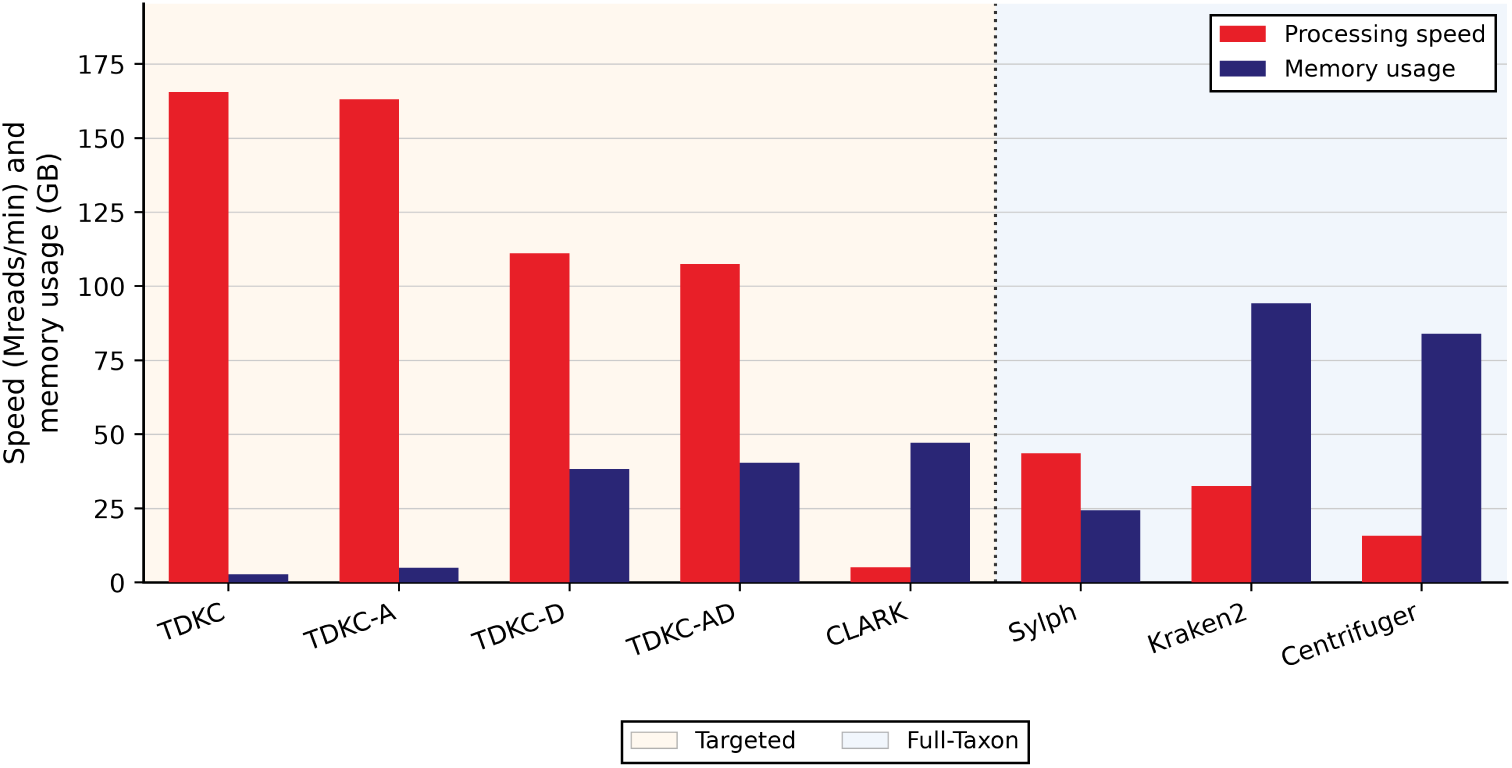
Performance comparison of TDKC against baseline classifiers (CLARK, Sylph, Centrifuger, Kraken2). TDKC was distilled on 68 target pathogen taxa with Homo sapiens and the bacteriophage MS2, and CLARK was built from the same set of accessions whose taxIDs fall under the same target pathogen taxa; Centrifuger, Sylph and Kraken2 were built from the full-taxon reference. All classifiers were queried using the same set of 96 clinical nasopharyngeal swab samples, a total of 494M reads. Processing speed is reported in millions of reads per minute, and memory usage is the peak resident set size (RSS). Full results are available in Table S1.

TDKC-A (with accession tracking) used 19.0× less memory than full-taxon Kraken2 (5.0 GB) and had nearly identical speed to TDKC (182s). TDKC-D (with domain-level detection) used 2.5× less memory (38.2 GB) and was 3.4× faster (267s) than the full-taxon Kraken2 database. With both features on, TDKC-AD introduced a minor overhead in both peak memory and speed to TDKC-D (Table S1).

### Evaluation framework

To determine target pathogens’ presence and absence in a sample, we evaluated all classifiers on contrived samples, which are known negative samples with synthetic material from a single virus added to them. Because we know exactly what is present, a TP is a read assigned to the spike-in virus, counted at the species level so that reads resolving to the spike-in species or to any node below it (e.g., a strain or isolate of that species) count toward the target. A FP is a read assigned to a different target on the panel or to any node below that target. Reads assigned to an ancestor of the spike-in species in the taxonomic lineage are treated as neither positive nor negative, as a less specific assignment to the correct organism is not a misclassification to a competing pathogen. *Homo sapiens* and the *bacteriophage MS2* are present in every sample by design and reads assigned to either are excluded from both TP and FP counts.

### TDKC produces minimal false positives while preserving sensitivity

To validate false-positive rates, we performed a limit-of-detection (LOD) experiment using the contrived spike-in samples described in Methods, with unspiked negative controls included to confirm the absence of background viral contamination. Reads were classified against the full-taxon database (Kraken2, Centrifuger, Sylph), CLARK (targeted), TDKC, and TDKC-D.

Across all spike-in and negative control samples, TDKC produced the fewest or tied-fewest FP reads among the per-read classifiers, and TDKC-D produced FP counts closely matching full-taxon Kraken2. Aggregated over 18 samples per spike-in virus, TDKC reduced FP reads by an average of 15 per virus relative to full-taxon Kraken2 (Table 1). On the six unspiked negative control samples, TDKC reported only 3 FP reads, while TDKC-D and full-taxon Kraken2 each reported 4, showing that TDKC’s low FP rate is maintained even in samples without any pathogen present. The TP read counts of these three tools were near-identical, agreeing within 0.65% per virus (Figure 4).

**Table 1.** False positive read counts across contrived LOD samples for targeted (TDKC, TDKC-D, CLARK) and full-taxon (Kraken2, Centrifuger, Sylph) classifiers. Total reads are reported in millions (M). FP counts are aggregated across all samples (N = 18 per virus; N = 6 for negative control).

| Target | Samples ($N$) | Total Reads | Targeted | | | Full-taxon | | |
| --- | --- | --- | --- | --- | --- | --- | --- | --- |
|  |  |  | TDKC | TDKC-D | CLARK | Kraken2 | Centrifuger | Sylph |
| SARS-CoV-2 | 18 | 81 M | 1 | 11 | 44,968 | 13 | 2,167 | 0 |
| Influenza A (H1N1) | 18 | 87 M | 3 | 12 | 37,610 | 12 | 1,583 | 0 |
| Influenza A (H3N2) | 18 | 78 M | 37 | 43 | 33,512 | 38 | 2,828 | 0 |
| Influenza B | 18 | 81 M | 437 | 448 | 46,241 | 491 | 3,170 | 0 |
| RSV | 18 | 84 M | 33 | 38 | 41,009 | 33 | 2,211 | 0 |
| Negative Control | 6 | 27 M | 3 | 4 | 11,545 | 4 | 260 | 0 |

**Fig. 4.**
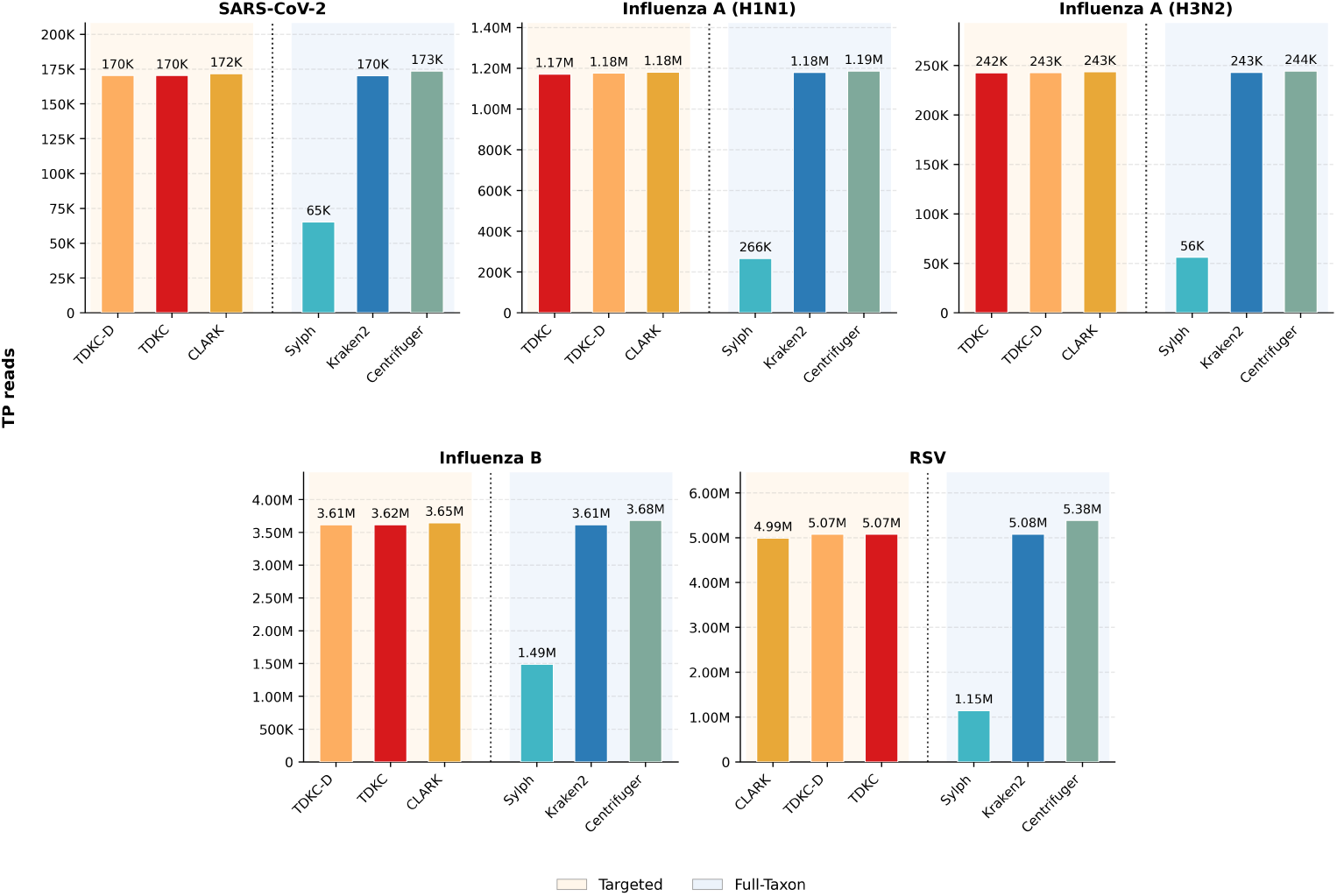
True positive (TP) read counts across classifiers for five spike-in viruses. Bars show aggregated TP read counts across N = 18 samples per virus for targeted classifiers (TDKC, TDKC-D, CLARK) and full-taxon classifiers (Kraken2, Centrifuger, Sylph). Sylph values are abundance-derived pseudo-counts (see Methods for details). Y-axes are scaled per panel to make small differences visible.

Centrifuger and CLARK produced substantially higher FP, and both showed elevated sensitivity relative to TDKC. Although Centrifuger draws on the same full-taxon reference as Kraken2, it produced 1,583 to 3,170 FP reads per virus over 18 samples, 7-fold to over 2,000-fold above TDKC, and reported 260 FP reads on the negative controls compared with 3 for TDKC (Table 1). CLARK was worse, with the highest FP of any classifier: 106-fold to nearly 45,000-fold more FP reads than TDKC, and 11,545 FP reads on the negative controls. These FP came alongside higher TP than TDKC: Centrifuger reported the highest TP of any classifier for all five viruses, from 0.7% to 6.1% above TDKC, while CLARK was only marginally higher for four of five viruses, by 0.4% to 0.9% (Figure 4). That Centrifuger and Kraken2 diverge so sharply in FP despite sharing the same reference indicates that FP control is governed by the classification algorithm rather than reference breadth alone. CLARK’s target-only index, which lacks the non-target filtering step that controls FPs, shows the same failure mode observed when Kraken2 is built from target sequences alone (Figure 2).

Sylph sat at the opposite extreme. It reported zero FP reads across all samples, but its TP read counts were only 23% to 41% of TDKC’s, consistent with sketch-based profiling trading sensitivity for low memory and speed.

Together, these results show that TDKC and TDKC-D keep FP minimal while preserving the pathogen signal, unlike the more sensitive tools that gained TP only at a substantial false-positive cost.

TDKC produced ambiguous classifications at consistently low rates across all samples (0.03% to 0.16% of total reads). In every sample, *>* 99% of ambiguous reads were ties involving either *Homo sapiens* or the MS2 spike-in control. The remaining tens of reads across all 96 contrived samples were pathogen-pathogen ties (e.g., Influenza A + Influenza B in Flu B spike-ins). This pattern may be consistent with chimeric reads introduced during sequencing, in which one mate of a paired-end read derives from a different source than the other.

## Discussion

TDKC allows researchers and clinicians to detect target pathogens in metagenomic samples faster and more memory-efficiently. In the base TDKC index, efficiency comes from discarding k-mers irrelevant to target pathogens, which results in a compact index. When domain-level detection is on (TDKC-D), the index retains the same volume of k-mer information as the full-taxon database, yet preserves efficiency by forgoing species-level classification outside of the target, which is not needed for domain-level detection by design.

TDKC’s speed comes from the architecture of the query pipeline. Decompression, classification and output run as independent stages connected by bounded channels. Dedicated reader threads decompress and parse input concurrently with classification. Worker threads take batches from a lock-free channel and accumulate results in thread-local buffers, acquiring no shared lock during classification. Lastly, a single writer absorbs the output for a final report. Because input handling is not serialized behind a shared lock, it can itself run in parallel. At high thread counts both tools are limited by their input stage, so this difference sets the throughput ratio. TDKC improves 13.7× from 1 to 32 threads, whereas Kraken2 improves 3.7× and saturates at approximately four-threads (Figure S1). Kraken2 performs decompression, FASTQ parsing and record copying inside a single OpenMP critical section, so only one thread can supply input at a time. Instrumentation shows its worker threads spend 91% of wall-clock time blocked on that section.

TDKC’s index size depends on the number of targets and their taxonomic rank. The index size does not necessarily grow linearly with the number of targets; it depends on how well-represented each target taxon is in the reference database and at what taxonomic rank it is defined. Heavily sequenced pathogens, such as HIV-1, Enterovirus, and Influenza A (Figure S2), have a large number of reference sequences and contribute more unique k-mers than uncommon pathogens. Additionally, target taxa placed at higher taxonomic ranks cover more descendant nodes, which results in larger clade-specific k-mer sets.

Importantly, TDKC and its different modes do not sacrifice accuracy for computational gain: they achieved nearly identical, and in some cases slightly better sensitivity and FP rates compared to full-taxon classification. For TDKC, the discrepancies in both FP and TP read counts are due to the presence of non-target taxa in the full-taxon database that are absent from TDKC’s index. In the full-taxon database, k-mers in a query read that match non-target organisms contribute additional hit groups, which can shift classification outcomes in either direction. When the non-target organism is taxonomically related to a target pathogen, its additional hit groups strengthen the target signal and can increase TP counts. On the other hand, when the non-target organism is unrelated, its hit groups may compete with and dilute the target signal, reducing TP counts. The same mechanism applies to FP: non-target hit groups can either push borderline reads over the classification threshold, increasing FPs, or introduce competing signals that suppress them. For TDKC-D, the source of discrepancies is slightly different. The index itself has the same amount of k-mer information as the full-taxon database, since it now stores non-target k-mers in Bloom filters. Because Bloom filters are probabilistic, they allow a small, bounded FP rate by design, and TDKC-D applies a penalty to domain hit groups to account for this. As a result, the same hit-group dynamics described above produce the small discrepancies in TP and FP read counts relative to the full-taxon database.

The accuracy of all classifiers ultimately depends on the quality of the full-taxon database. Theoretically, target k-mers in TDKC’s distilled index should carry the same LCA-resolved taxIDs as Kraken2’s, assuming no minimizer collisions. This means that sensitivity depends on how well each target pathogen is represented in the reference. If a target is poorly represented and the circulating strain in a sample is novel, both the full-taxon database and TDKC may fail to detect the pathogenic signal even when it is present. Moreover, k-mers that are extracted from contaminated or mislabeled reference sequences in the full-taxon database will be distilled directly into TDKC’s index, keeping those errors. This is reflected in our results (Table 1), where the full-taxon database itself produces FPs, and TDKC is concordant with them. TDKC’s detection performance depends heavily on how clean and well-curated the reference sequences in the full-taxon database are.

## Conclusions

TDKC reframes metagenomic classification for the diagnostic setting, where the goal is to detect a defined panel of pathogens rather than profile an entire sample. By distilling only target-relevant k-mers from a full-taxon reference and removing those shared with non-target organisms, TDKC reduces memory by 16.9–33.6× and classification runtime by 5.1–34.7× relative to per-read classifiers, while preserving the sensitivity and false-positive control of full-taxon Kraken2. Accession tracking and domain-level detection extend its utility toward subtype analysis and broad taxonomic screening without sacrificing efficiency. As reference databases and sequencing throughput continue to grow, targeted distillation offers a practical path to scaling pathogen detection across thousands of clinical samples and large sequence repositories.

## Methods

### Datasets

#### Reference sequences for index construction

The full-taxon reference was constructed from NCBI RefSeq release 230, comprising bacterial, archaeal, viral, human (*Homo sapiens*), plasmid, and UniVec Core sequences. To capture circulating strains absent from curated repositories, it was supplemented with Viral NT sequences from the NCBI Nucleotide database, yielding 4,679,153 accessions in total. All reference sequences were low-complexity filtered with *dustmasker*[28] before index construction.

From this reference we defined a target set of 68 pathogen taxa representing the most common and high-consequence human pathogens, with emphasis on respiratory and enteric viruses, together with *Homo sapiens* and the *bacteriophage MS2* as a spike-in control (Table S2). The target accessions are the subset of the full reference whose taxIDs fall under any target taxon or its descendants. To ensure a fair comparison, all targeted classifiers (TDKC and CLARK) were built from this identical target-accession subset, and all full-taxon classifiers (Kraken2, Centrifuger, Sylph) were built from the complete reference described above.

#### Query datasets

Benchmarking used two sets of query reads. Accuracy was evaluated on contrived LOD samples, where ground truth is known. These were prepared from ZeptoMetrix reference materials for five respiratory viruses (Influenza A H3N2, SARS-CoV-2, Influenza A H1N1, Influenza B, and RSV-A), each spiked into a negative nasal swab matrix at six serial 1:5 dilutions in triplicate (N = 18 per virus), alongside six unspiked negative-control samples. Computational performance was evaluated on a clinical cohort of 96 multiplexed nasopharyngeal swab samples from a single sequencing run (Illumina NextSeq 2000, 2×51 bp), totaling 494 million reads. Reads were input into all classifiers as raw, gzipped FASTQ without quality trimming, adapter removal, or host filtering.

### Target-specific k-mer distillation from reference sequences

The distillation process constructs a target-specific minimizer index directly from reference sequences, without requiring a pre-built database. The input consists of a reference FASTA file, the NCBI taxonomy, and a user-defined list of target taxIDs.

A naive approach would extract minimizers from every sequence in the full reference FASTA, compute LCA assignments across all of them, and then keep only those whose final LCA falls within a target clade. This is how a standard Kraken2 database build operates, and it requires tracking the taxonomy of every minimizer across the entire reference, which is both memory-intensive and slow. Instead, TDKC exploits a key observation: the set of minimizers extracted from target sequences alone is a superset of the final target-distilled index. Some of these minimizers will ultimately be excluded because they also appear in non-target sequences. This motivates a two-phase design. In Phase 1, TDKC extracts minimizers and resolves their taxonomy using only target sequences, eliminating the need to track billions of irrelevant minimizers. In Phase 2, TDKC removes any target minimizers that also appear in non-target sequences.

To prepare for Phase 1, TDKC extracts accession headers from the FASTA headers and maps each accession to its corresponding taxID using the NCBI accession2taxid files. Accessions whose taxIDs fall within any target taxon or its descendants are identified as target accessions, and their sequences are extracted into a target sub-FASTA. This reduces the data processed in Phase 1 from the full reference (often hundreds of gigabytes) to only the target-relevant subset.

#### Phase 1: Target minimizer extraction

TDKC extracts minimizers from each target sequence using a k-mer length of *k* = 35, a minimizer length of *ℓ* = 31, and a spaced seed mask with *s* = 7 positions masked. Spaced seed masking is applied to tolerate mismatches and increase sensitivity. These settings match the default parameters of Kraken2, whose parameter sweep over *k, ℓ* and *s* identified this configuration as providing a favorable balance of accuracy, classification speed, and memory usage[26]. Each minimizer is assigned the taxID of its source accession. When the same minimizer is encountered across multiple target accessions, its taxonomy is resolved by computing their LCA. After processing all target sequences, each minimizer carries the LCA of all target accessions in which it appeared. Minimizers whose LCA falls outside the relevant target clades are discarded.

Targets can be defined at any taxonomic rank depending on the desired level of detection. For broad detection, a user might specify a genus-level taxID (e.g., Enterovirus); for more specific detection, a species-level taxID (e.g., Rhinovirus A). Because target accessions include sequences from the target taxon itself and all of its descendants, the minimizers extracted from those accessions span multiple taxa below the user-specified target. To ensure these descendant-level minimizers still contribute to detection of the user’s target, TDKC applies a “roll-up” during label assignment. Each minimizer climbs the taxonomy tree to its nearest target ancestor and is reported under that target rather than its own finer-grained taxID (Figure 1a). When nested targets are specified – for example, both a genus and a species within that genus – minimizers inside the species halt at the species, and minimizers elsewhere in the genus halt at the genus.

#### Phase 2: Non-target challenge

The target minimizer set from Phase 1 may still contain minimizers shared with non-target organisms. If retained, these shared minimizers would produce false-positive classifications during the query phase. We refer to the filtering step as the “non-target challenge”: non-target sequences challenge each target minimizer, and any minimizer found in both target and non-target sequences is dropped from the index. TDKC performs this by scanning all non-target sequences in the full reference FASTA, extracting their minimizers, and testing each for membership in the target set. Any matching minimizer is removed from the target set (see Figure S3 for detailed retention numbers at each stage). Importantly, this phase requires no expensive LCA computation or taxonomy tracking. If a target minimizer also occurs in a non-target sequence, computing its LCA would push it outside the target clade, since the non-target node lies outside by definition. A simple membership test therefore suffices. This makes the challenge phase significantly faster than a full taxonomic resolution over the entire reference (Figure S4). Target accessions are skipped during this scan since they were already processed in Phase 1.

### Compact k-mer indexing with minimal perfect hashing

To minimize memory footprint, TDKC indexes its target minimizers using a Minimal Perfect Hash Function (MPHF), constructed with BBHash[29]. An MPHF maps N keys to the integer range [0,N-1] without storing the keys themselves, eliminating the memory overhead of a conventional hash table. Because the keys are not retained, additional structures are needed to reject out-of-set queries and associate each minimizer with its taxonomic label. TDKC maintains two parallel arrays indexed by the MPHF output: a 16-bit fingerprint array (.fp), which stores a truncated hash of each minimizer for membership verification, and a local taxID index array (.taxid), which stores an 8-bit internal index for each minimizer.

To resolve this internal index to an NCBI taxID, TDKC maintains a separate taxID lookup table (.taxmap) that maps each internal index to its corresponding 32-bit NCBI taxID. This is a deliberate memory optimization: NCBI taxIDs are 32-bit integers, but the number of distinct target taxa is typically small (e.g., 70 in our default configuration). By storing a compact 8-bit internal index per minimizer and resolving it to the full NCBI taxID only at report time, TDKC reduces the per-minimizer taxonomy overhead from 4 bytes to 1 byte, which is a 4-fold savings on the .taxid array.

At query time (Figure 5a), a minimizer extracted from a read is passed through the MPHF to yield an index i. TDKC then compares the minimizer’s fingerprint against .fp[i]: if they match, the minimizer is assumed to be a member of the index, and its taxID is retrieved by looking up .taxid[i] in .taxmap. Theoretical collision rate for a minimizer that is not present in the MPHF is bounded by the probability of a random 16-bit hash collision, which is 1*/*2^16^ (≈ 0.0015%).

**Fig. 5.**
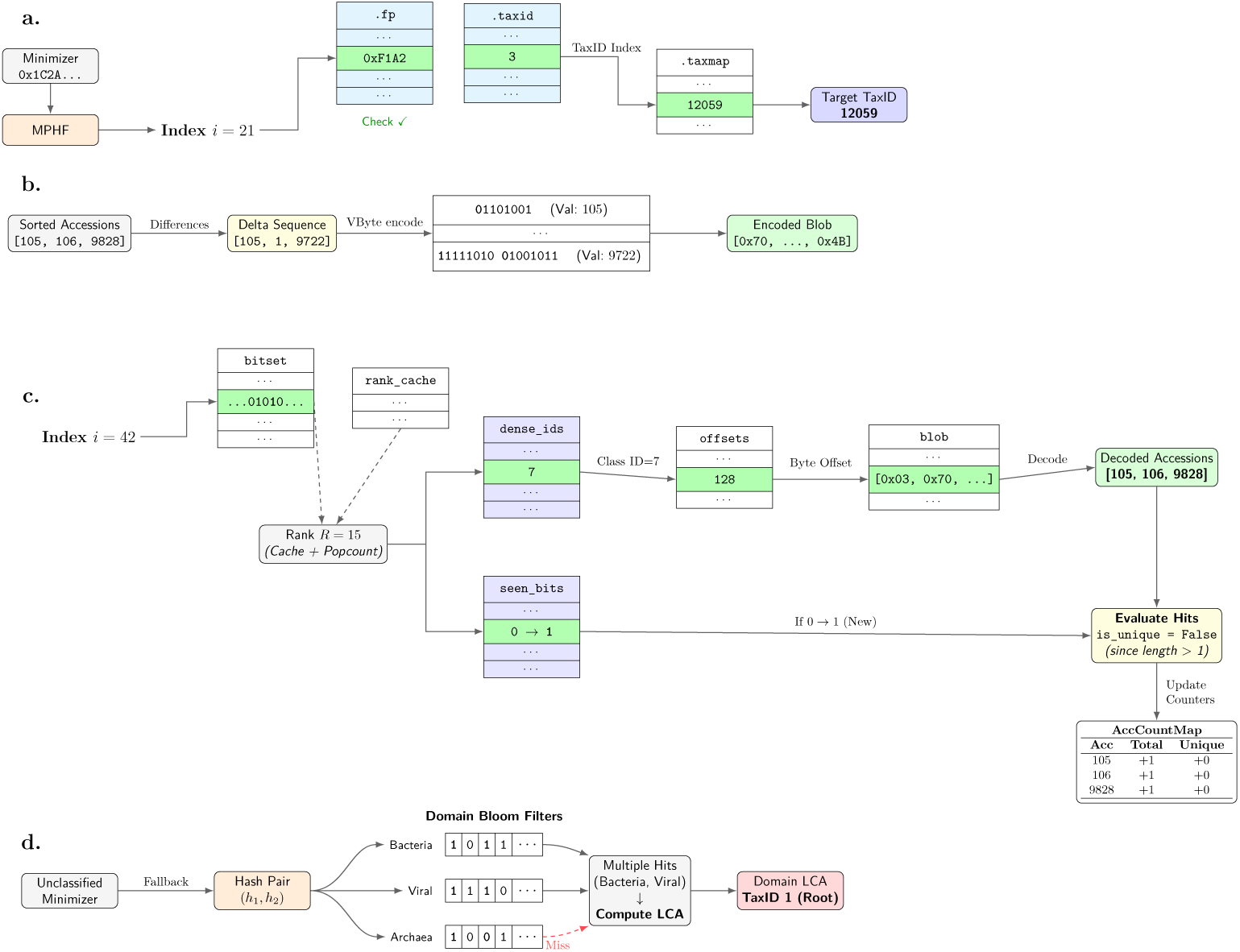
Database architecture and memory management. Within each subpanel, arrays of the same color indicate identical length. (a) The lookup mechanism maps a minimizer to a taxID index via a Minimal Perfect Hash Function (MPHF), verified against a fingerprint array. (b) Delta and Variable-Byte (VByte) encoding compresses sorted accession IDs. (c) The succinct data structure resolves a minimizer index into an equivalence class, decoding it into variable-length accession IDs and updating the accession count map. (d) Unclassified minimizers are queried against Bloom filters for each domain to perform domain level classification. Hits across multiple domains are resolved by computing the LCA.

While the per-minimizer fingerprint collision rate is small, spurious hits could in principle accumulate across the minimizers in a read. For a 2×51 bp paired-end read with *k* = 35, *ℓ* = 31, each mate contributes *L* − *k* + 1 = 17 k-mers, so a read pair contains *n* = 34 k-mers. Under the standard minimizer-density approximation, the expected number of distinct minimizers per read pair is roughly 2*n/*(*k* − *ℓ* + 2) ≈ 11. The probability that two of the minimizers collide with indexed entries is at most (11/2) · (2^−16^)^2^ ≈ 10^−8^ per read pair, or ∼0.06 spurious classifications per 5M read pair sample. Because TDKC requires at least two distinct minimizer hit groups for any classification (whether resolved to a single target or reported as ambiguous), both bounds apply equally to false-positive and ambiguous outputs. These predicted rates are much smaller than the FP and ambiguous read counts observed in the evaluation of contrived samples, confirming that MPHF fingerprint collisions are not a meaningful source of either FPs or ambiguous classifications.

### Memory-efficient accession tracking via succinct encoding

Tracking the set of accessions from which each target minimizer originated is useful for subtype classification and quantification, as well as for debugging FP minimizer origins. However, naively storing per-minimizer accession lists is expensive in memory, because many accessions produce the same minimizer.

To address this, we group minimizers that share an identical set of source accessions into equivalence classes. Each class stores its accession list once, and all minimizers in that class reference only the shared class ID. Within each class, the sorted accession IDs are delta-encoded; this stores only the difference between consecutive IDs. Then deltas are compressed with Variable-Byte (VByte) encoding. This essentially encodes integers in fewer bytes by using 7-bits per byte for the payload and 1-bit as a continuation flag (Figure 5b). All encoded lists are concatenated into a single contiguous byte blob, with a separate offset array recording where each class begins. To resolve a minimizer to its accession list at query time, we build a bit vector over the full target minimizer index space. A set bit indicates that the corresponding minimizer carries accession data. Not all minimizers require tracking. By default, human minimizers are excluded to save memory and users may specify to exclude additional taxIDs. Given a minimizer index, a query on this bit vector yields the position in a dense array of equivalence class IDs. This takes O(1) with the help of a precomputed pop-count cache (Figure 5c).

To maximize delta encoding compression and make database building deterministic, accession IDs are presorted by taxonomic lineage. For each accession, we trace its taxID upward through the NCBI taxonomy tree to the root, producing a lineage path as a sequence of node IDs. Accessions are then sorted lexicographically by lineage, which groups biologically related sequences adjacent to each other. This ultimately yields smaller deltas and better compression. Because the sort is deterministic, the database build is also deterministic regardless of the sequence order in the FASTA file

Together, these design choices yield substantial compression. In our target configuration, accession tracking is applied to 38.5M minimizers out of 0.84B in the index, since excluded human minimizers dominate the index by volume (Figure S5b). These tracked minimizers collapse into 9.8M equivalence classes (Figure S5a), and lineage-sorting produces highly compressible deltas: 63.1% encode in a single byte and 95.5% in two bytes or fewer (Figure S5c). Combined, these optimizations reduce the accession index from 8.2GB under a naive per-minimizer u32 layout to 1.9GB, a 4.3-fold reduction (Figure S5d).

When accession tracking is enabled, TDKC-A also outputs a strain report alongside the per-read output. For each distinct minimizer hit in the sample, TDKC-A resolves it to an equivalence class through the dense array lookup described above and increments a total-hit counter for each member accession, plus a unique-hit counter when the class has size one. (Figure 5c) To prevent repetitive minimizers from inflating these counts, each distinct minimizer is counted at most once per sample, enforced by an atomic seen_bits bitset. The total-hit counter therefore reflects the per-accession minimizer coverage from the sample, while the unique-hit counter isolates the subset of that coverage unique to a single accession, providing a strain-specific signal.

### Efficient domain-level detection via probabilistic Bloom filters

To support domain-level classification (e.g., bacteria, archaea, and viruses) without the memory overhead of a full-taxon hash table, TDKC uses Bloom filters.

During the build phase (TDKC-D), TDKC performs a two-pass scan over reference sequences of each domain. The first pass uses the HyperLogLog++[30, 31] algorithm to estimate the total cardinality of unique minimizers. This estimate is used to dynamically allocate the exact number of bits and hash functions (*h*) required for the filter to achieve a user-defined FP rate. Because metagenomic classification requires querying millions of reads, the default Bloom filter FP rate is set to 0.01% to minimize FP hits. However, users may increase this to 0.1% if approximate profiling is sufficient.

Because computing *h* independent hash functions is computationally inefficient, TDKC instead uses double-hashing technique, where two hash functions are combined to simulate many hash functions while maintaining a good distribution of bits[32]. In the second pass, TDKC extracts minimizers from the sequences and inserts them into the filter. During the query phase, any minimizers in a read that fail to pass the MPHF and fingerprint checks are subsequently queried against the loaded domain Bloom filters. If a minimizer matches multiple domain filters, TDKC resolves it using a precomputed cross-domain LCA: bacteria and archaea hits collapse to “cellular organisms” (taxID 131567), while any combination involving viral hits collapses to “root” (taxID 1)(Figure 5d).

### Read classification

For a read to be classified, TDKC requires a minimum number of distinct minimizer hit groups as in Kraken2. The default threshold is set to 2. When domain-level detection is on, TDKC calculates its effective hit groups using the following formula:

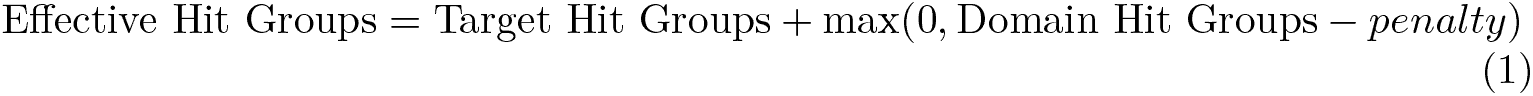

This applies a penalty (default: 1) to domain hit groups to account for the probabilistic nature of the Bloom filters and to ensure that broad domain evidence does not disproportionately influence the classification of borderline reads. For paired-end reads, hit groups are simply concatenated across both mates. When a read’s effective hit group count meets the required threshold, each matching target and domain is treated as a valid target candidate. TDKC computes a path weight for each candidate by summing the candidate’s direct hits and the hits of all its ancestors, which incorporates evidence from higher taxonomic nodes. Finally, the read is classified as the candidate with the maximum path weight. To avoid node-by-node traversal of a full taxonomy tree, TDKC uses a precomputed *N* × *N* Boolean ancestor matrix (where N is the number of target taxa) that encodes ancestor-descendant relationships between all pairs.

If targets are tied, TDKC resolves the tie by computing the LCA. When domain-level detection is on, near-root taxa (root, Bacteria, Archaea, Viruses, cellular organisms) are eligible LCA candidates, so tied reads are guaranteed to resolve to a near-root ancestor. When domain-level detection is off, these near-root taxa are excluded as LCA candidates, and if no common ancestor exists among the tied targets, the read is classified as “ambiguous”. When accession-level tracking is on, TDKC-A also reports a set of source accession IDs resolved from its equivalence class.

### Benchmarking

We compared TDKC against Kraken2 (v2.1.3), Centrifuger (v1.1.2), and Sylph (v0.9.0) for full-taxon classification and CLARK (v1.3.0.0) for targeted classification. Targeted classifiers (TDKC, CLARK) were built from the target-accession subset and full-taxon classifiers (Kraken2, Centrifuger, Sylph) from the complete reference, so that all tools indexed the same underlying sequences.

Kraken2 and CLARK databases were built with default parameters. Centrifuger was built with –build-mem 380G. Sylph reference sketches were built with subsam-pling rate -c 100. For classification, Kraken2, Centrifuger, and CLARK were run with default parameters; Sylph was run with –min-number-kmers 10.

Because Sylph does not make a per-read classification, we ran a custom script that maps each profiled accession to its NCBI taxID, aggregates Sylph’s sequence-abundance estimates along the taxonomic lineage, and converts each taxon’s relative abundance into a pseudo-read count by scaling it to the sample’s total read count.

All query benchmarks, and all index builds except Centrifuger’s, were run on an Intel Xeon Platinum 8175M (2.50 GHz) node with 128 GB RAM using 32 threads. Centrifuger’s index build required 363 GB of peak memory, exceeding the capacity of this node, and was therefore computed on a separate Intel Xeon E5-2650 v4 node with 503.8 GB RAM using 23 threads. Centrifuger’s build time (Figure S4) is not directly comparable to those of the other tools.

Classification times are wall-clock time measured after the index was loaded. Index load times are reported separately. Each value is the mean of three runs. For each tool, the index was loaded once per job and samples were processed sequentially. Peak memory was measured using the command /usr/bin/time -v.

## Declarations

### Ethics approval and consent to participate

The use of clinical swab and contrived samples in this study was approved by the UCLA Institutional Review Boards (studies IRB-22-1890 and IRB-25-2422).

### Consent for publication

Not applicable.

### Availability of data and materials

TDKC is implemented in Rust and available at https://github.com/543090lee/TDKC, and archived in Zenodo under an MIT license [33]. The target-distilled TDKC database used in this study is available in Zenodo under a CC BY 4.0 license [34]. It was distilled from NCBI RefSeq release 230 [35] and Viral NT sequences from the NCBI Nucleotide database [36], both publicly available. All scripts used to generate the figures and analyses in this study are available in the project repository [37].

## Acknowledgements

Not applicable.

## Abbreviations

RefSeq: NCBI reference sequence database
TP/FP: True positives / False positives
TDKC: Target Distilled K-mer Classifier
LCA: Least common ancestor
taxID: NCBI taxonomy ID
MPHF: Minimal perfect hash function
RSS: Resident set size VByte: Variable-byte
LOD: Limit-of-detection

## Supplementary Materials

**Table S1.**
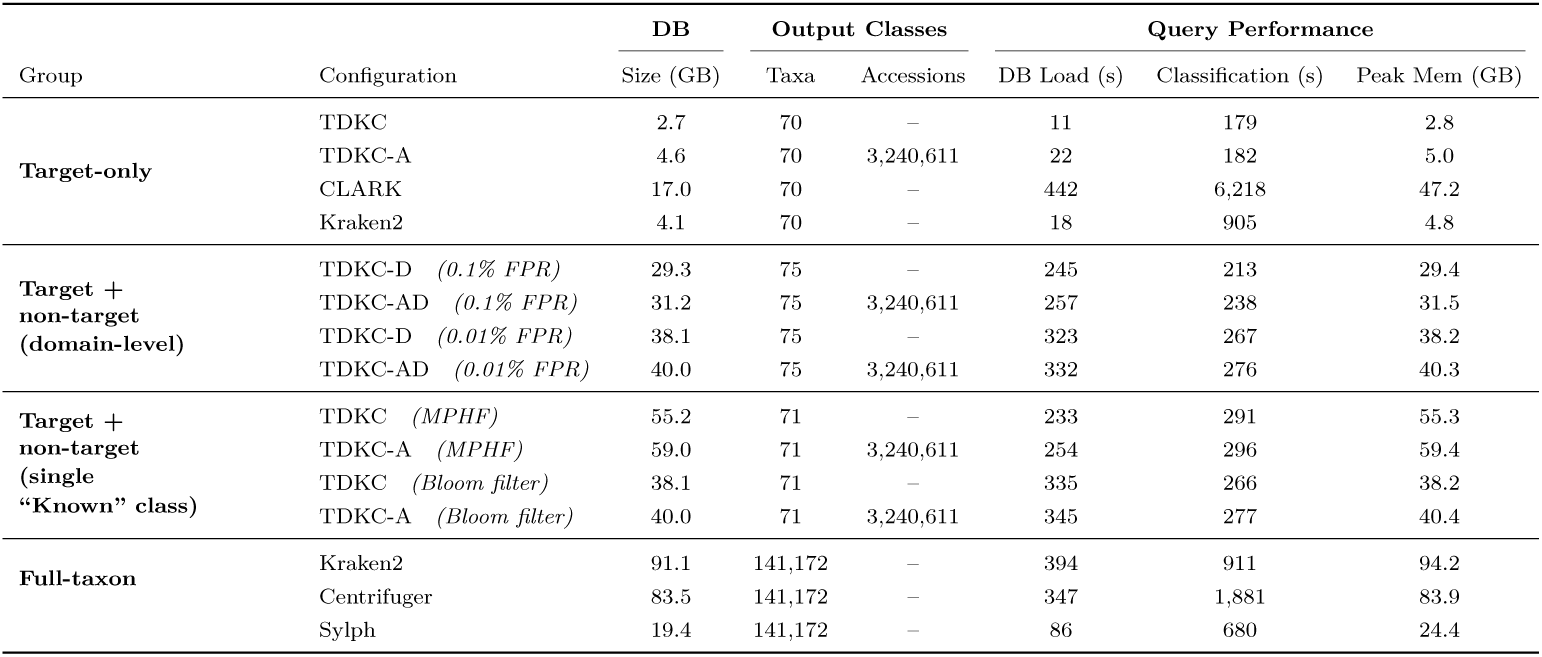
Database specifications and query performance of TDKC compared to baseline classifiers (Kraken2, CLARK, Centrifuger, Sylph). Target-only configurations index only the k-mers extracted from accessions whose taxIDs fall under the 68 target pathogen taxa; TDKC, CLARK, and target-only Kraken2 were all built from the same set of target accessions. All other TDKC configurations are built upon the same full reference sequences but differ in how non-target k-mers are stored and queried. Domain-level Bloom filters assign non-target k-mers to taxonomic domains (Bacteria, Archaea, Viruses) at different FP rates. Single “Known” class configurations collapse all non-target k-mers into one label, stored either in a Bloom filter (0.01% FP rate) or directly in the MPHF with target k-mers. Output classes show the number of distinct labels each method can assign to a k-mer. Accession tracking is available for targets. Domain configurations include 70 targets, 3 domain classes (Bacteria, Archaea, Viruses), and 2 ancestor nodes (cellular organisms, root) from cross-domain LCA resolution. Experimental conditions as in Figure 3.

| Group | Configuration | DB | Output Classes |  | Query Performance |  |  |
| --- | --- | --- | --- | --- | --- | --- | --- |
|  |  | Size (GB) | Taxa | Accessions | DB Load (s) | Classification (s) | Peak Mem (GB) |
| Target-only | TDKC | 2.7 | 70 | – | 11 | 179 | 2.8 |
|  | TDKC-A | 4.6 | 70 | 3,240,611 | 22 | 182 | 5.0 |
|  | CLARK | 17.0 | 70 | – | 442 | 6,218 | 47.2 |
|  | Kraken2 | 4.1 | 70 | – | 18 | 905 | 4.8 |
| Target +<br>non-target<br>(domain-level) | TDKC-D (0.1% FPR) | 29.3 | 75 | – | 245 | 213 | 29.4 |
|  | TDKC-AD (0.1% FPR) | 31.2 | 75 | 3,240,611 | 257 | 238 | 31.5 |
|  | TDKC-D (0.01% FPR) | 38.1 | 75 | – | 323 | 267 | 38.2 |
|  | TDKC-AD (0.01% FPR) | 40.0 | 75 | 3,240,611 | 332 | 276 | 40.3 |
| Target +<br>non-target<br>(single<br>“Known” class) | TDKC (MPHF) | 55.2 | 71 | – | 233 | 291 | 55.3 |
|  | TDKC-A (MPHF) | 59.0 | 71 | 3,240,611 | 254 | 296 | 59.4 |
|  | TDKC (Bloom filter) | 38.1 | 71 | – | 335 | 266 | 38.2 |
|  | TDKC-A (Bloom filter) | 40.0 | 71 | 3,240,611 | 345 | 277 | 40.4 |
| Full-taxon | Kraken2 | 91.1 | 141,172 | – | 394 | 911 | 94.2 |
|  | Centrifuger | 83.5 | 141,172 | – | 347 | 1,881 | 83.9 |
|  | Sylph | 19.4 | 141,172 | – | 86 | 680 | 24.4 |

**Table S2.** Default target pathogen taxID list: 68 pathogen taxa plus Homo sapiens and bacteriophage MS2.

| TaxID | Name | TaxID | Name |
| --- | --- | --- | --- |
| 290028 | Human coronavirus HKU1 | 10372 | Human betaherpesvirus 7 |
| 31631 | Human coronavirus OC43 | 689403 | Human bocavirus 1 |
| 277944 | Human coronavirus NL63 | 1335626 | Middle East respiratory syndrome-related coronavirus |
| 11137 | Human coronavirus 229E | 694009 | Severe acute respiratory syndrome-related coronavirus |
| 162145 | human metapneumovirus | 142786 | Norovirus |
| 12059 | Enterovirus | 208726 | Human hepatitis A virus |
| 42789 | enterovirus D68 | 28883 | Human parechovirus |
| 147711 | Rhinovirus A | 11676 | Human immunodeficiency virus 1 |
| 147712 | Rhinovirus B | 11709 | Human immunodeficiency virus 2 |
| 463676 | Rhinovirus C | 10407 | Hepatitis B virus |
| 11250 | human respiratory syncytial virus | 11103 | Hepatitis C virus |
| 12814 | Respiratory syncytial virus | 12461 | Hepatitis E virus |
| 11320 | Influenza A virus | 333760 | Human papillomavirus 16 |
| 11520 | Influenza B virus | 10798 | Human parvovirus B19 |
| 11552 | Influenza C virus | 11041 | Rubella virus |
| 114727 | H1N1 subtype | 1980442 | Orthohantavirus |
| 119210 | H3N2 subtype | 12637 | Dengue virus |
| 102793 | H5N1 subtype | 64320 | Zika virus |
| 2697049 | Severe acute respiratory syndrome coronavirus 2 | 186538 | Zaire ebolavirus |
| 12730 | Human respirovirus 1 | 3052225 | Henipavirus nipahense |
| 2560525 | Human orthorubulavirus 2 | 11292 | Lyssavirus rabies |
| 11216 | Human respirovirus 3 | 3052310 | Mammarenavirus lassaense |
| 11224 | Human parainfluenza virus 4a | 186537 | Marburgvirus |
| 10509 | Mastadenovirus | 3052223 | Henipavirus hendraense |
| 10359 | Human betaherpesvirus 5 | 37124 | Chikungunya virus |
| 10298 | Human alphaherpesvirus 1 | 11082 | West Nile virus |
| 11234 | Measles morbillivirus | 11089 | Yellow fever virus |
| 2560602 | Mumps orthorubulavirus | 11072 | Japanese encephalitis virus |
| 10912 | Rotavirus | 11588 | Rift Valley fever virus |
| 10244 | Monkeypox virus | 3052518 | Orthonairovirus haemorrhagiae |
| 10376 | human gammaherpesvirus 4 | 10255 | Variola virus |
| 10335 | Human alphaherpesvirus 3 | 9606 | Homo sapiens |
| 10310 | Human alphaherpesvirus 2 | 3418604 | Betacoronavirus pandemicum |
| 37296 | Human gammaherpesvirus 8 | 2509511 | Sarbecovirus |
| 12022 | Escherichia phage MS2 | 10368 | Human betaherpesvirus 6 |

**Fig. S1.**
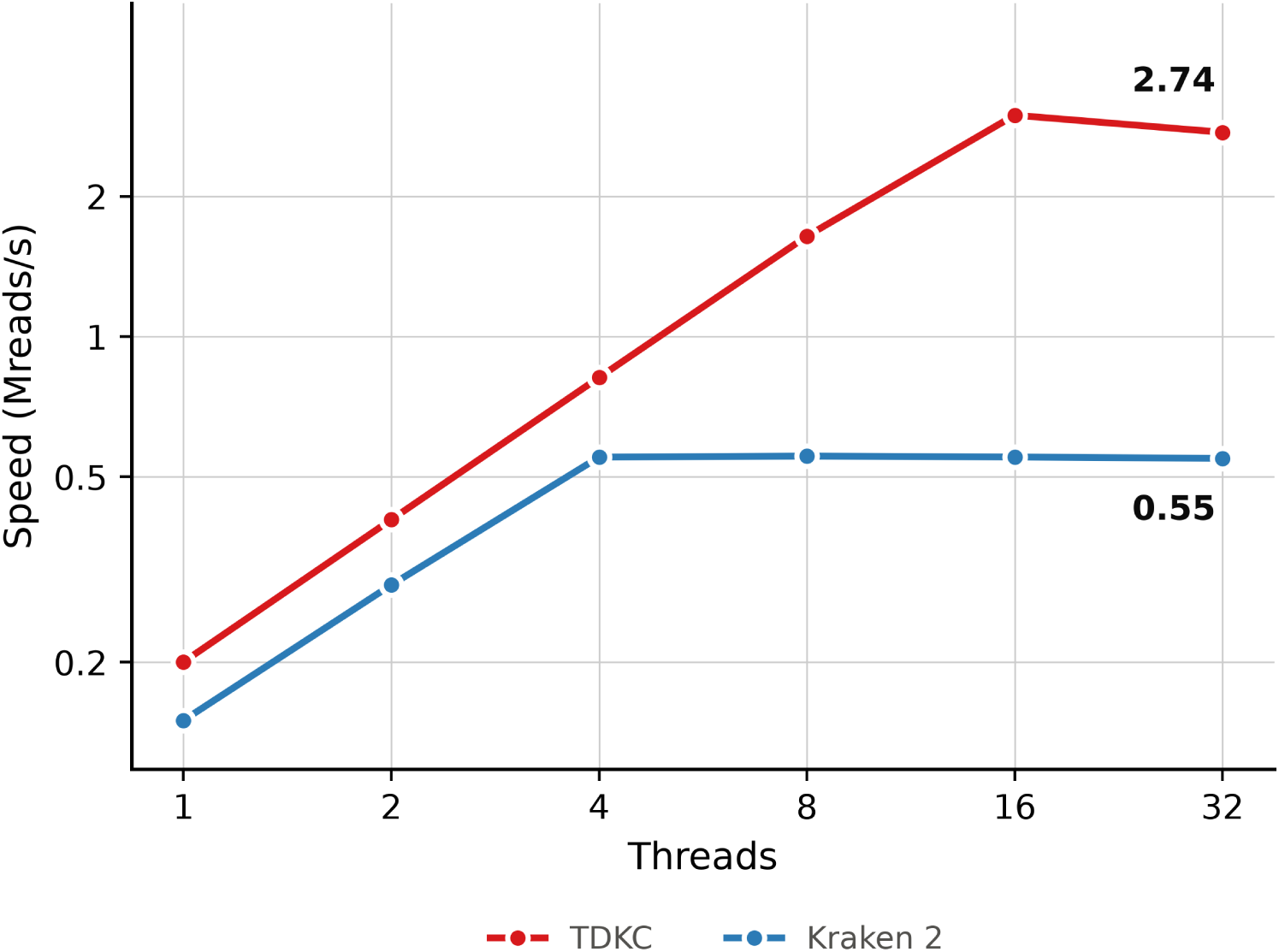
Thread scalability of TDKC and Kraken 2. Classification speed against thread count for a single clinical nasopharyngeal swab sample (5,755,242 paired-end reads). Kraken 2 saturates at four threads and holds 0.55 Mreads/s from 4 to 32 threads. TDKC continues to gain to 16 threads, reaching 2.99 Mreads/s, and delivers 2.74 Mreads/s at 32 threads. The gap widens with thread count, from 1.3× at one thread to 5.0× at 32.

**Fig. S2.**
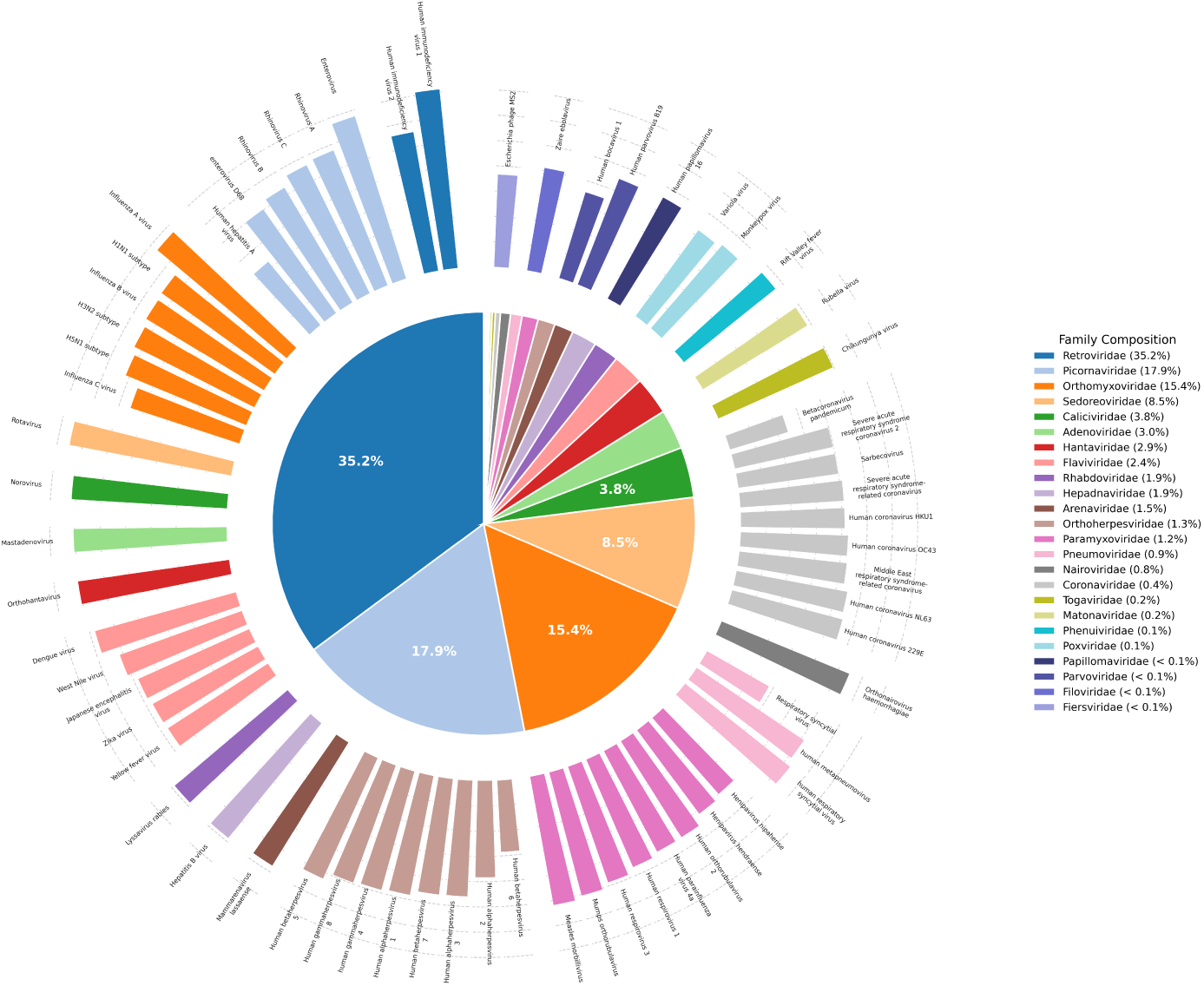
Human pathogen composition of the distilled database. Inner pie chart shows the proportion of target minimizers contributed by each viral family (Homo sapiens excluded). Outer ring shows the minimizer counts for each target taxonomy on a log10 scale. Both rings share the same family color scheme.

**Fig. S3.**
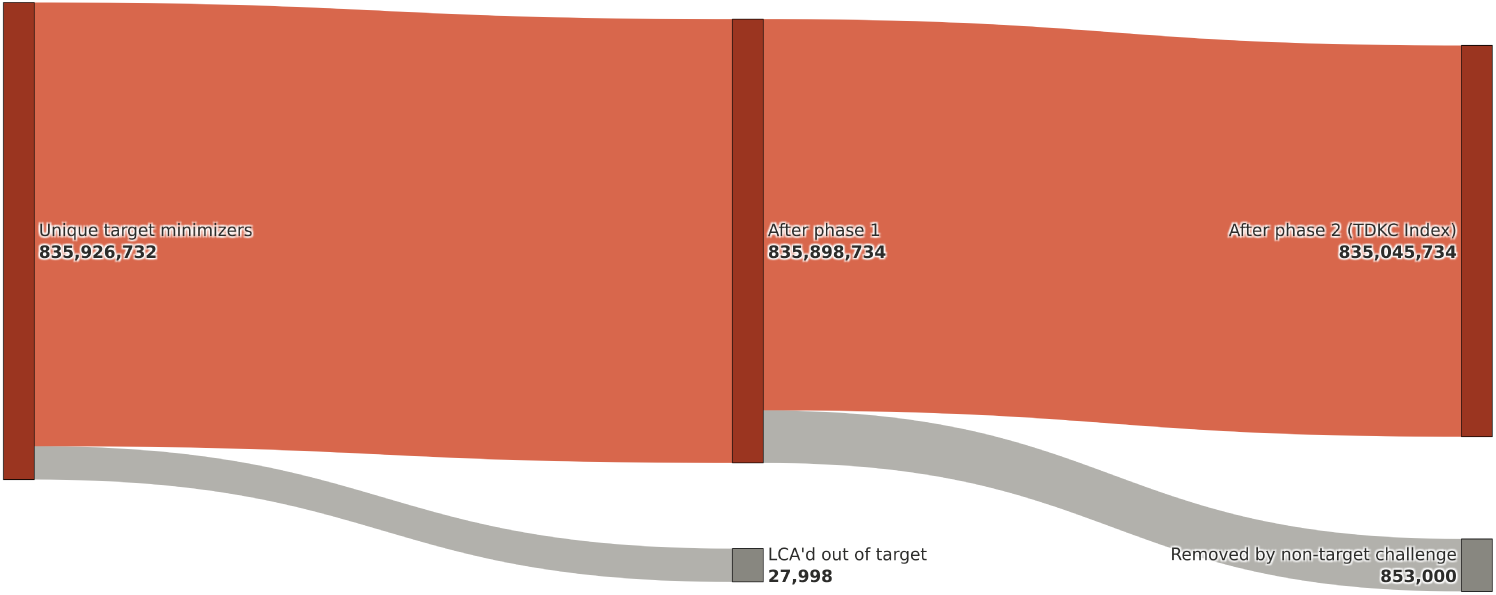
Number of unique minimizers retained at each stage of distillation. Phase 1 removed 27,998 minimizers whose LCAs fell outside the target clades. Phase 2 challenged the remaining target minimizers against non-target sequences, removing an additional 853,000 shared minimizers to produce the final index.

**Fig. S4.**
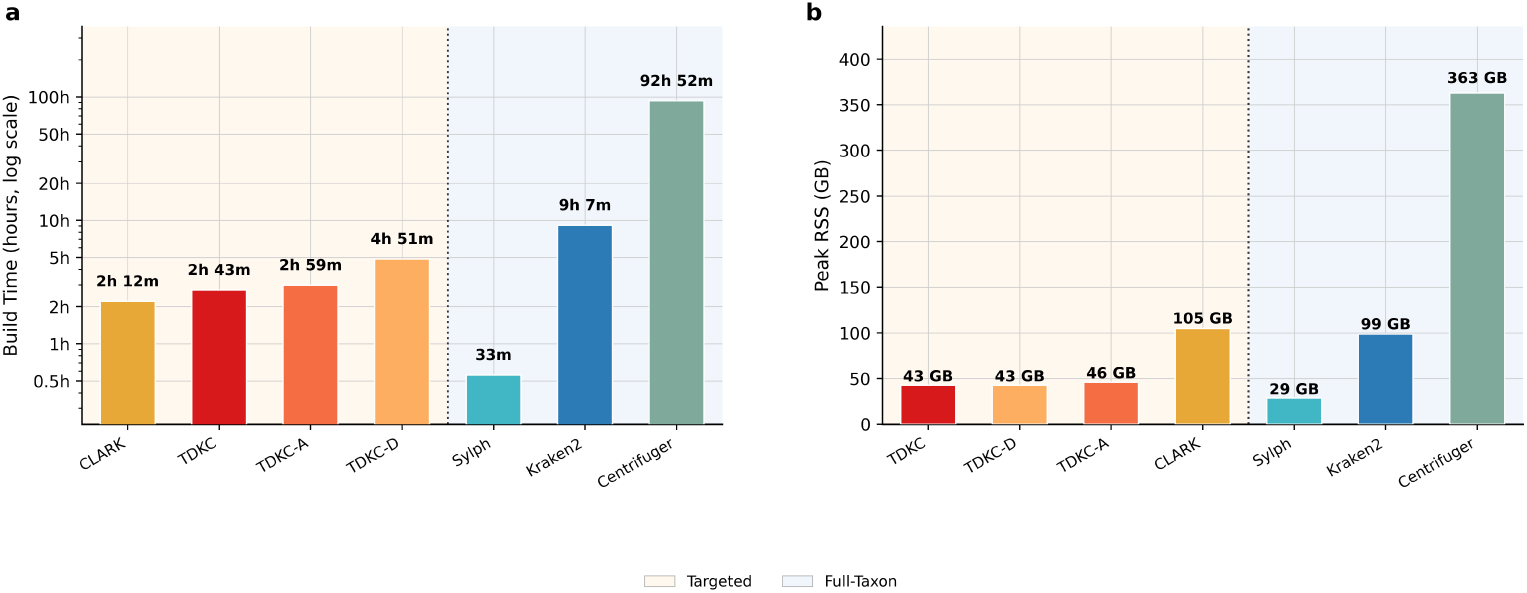
Database build time and peak memory usage. (a) Build time and (b) peak RSS for TDKC, TDKC-A, TDKC-D, and the baseline classifiers Kraken2, CLARK, Sylph, and Centrifuger. All builds used 32 threads except Centrifuger, which used 23 threads on a separate higher-memory node (see Methods for details).

**Fig. S5.**
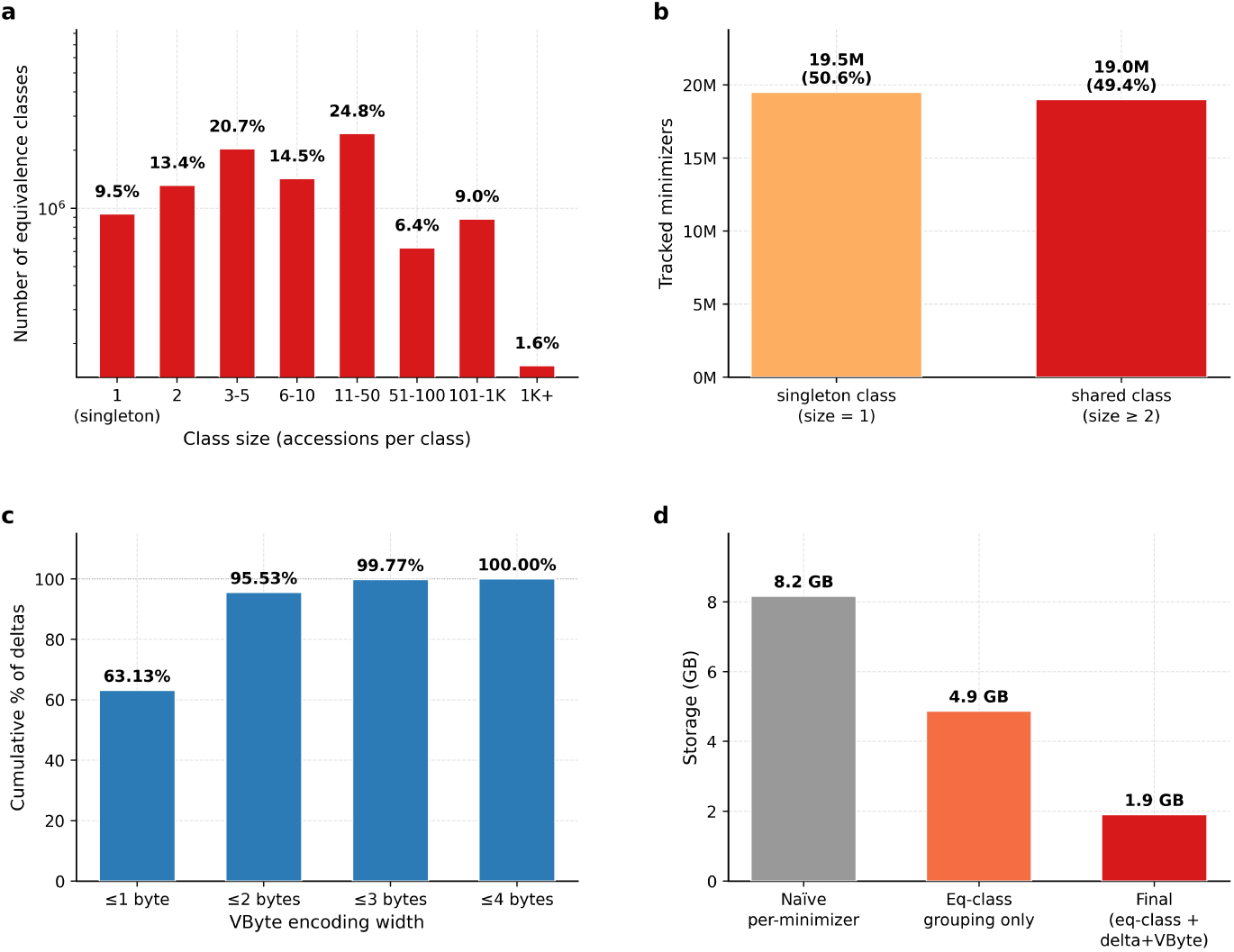
Accession database structure and storage efficiency in TDKC-A. (a) Distribution of equivalence-class sizes across 9.8M classes. (b) Of 38.5M minimizers with accessions tracked, half of them fall in singleton (size-1) classes. (c) Cumulative fraction of delta values encodable in 1, 2, 3 and 4 VBytes after lineage-sorting. 95.5% of deltas fit in ≤ 2 bytes (d) Accession index size under three encoding schemes: naive per-minimizer u32 lists (8.2GB), equivalence-class grouping with u32 (4.9GB), and the final encoding combining equivalence-class with delta+VByte compression (1.9GB). Final encoding results in a 4.3× reduction from the naive baseline.

